# Honeycomb cell walls are plastic; initially built as curves the walls are reformed by bees during subsequent construction

**DOI:** 10.1101/2022.07.13.499871

**Authors:** Vincent Gallo, Alice Bridges, Joseph Woodgate, Lars Chittka

**Author notes:** Corresponding author: Vince Gallo.

## Abstract

Honeybee comb comprises recognisable hexagons, each with straight sides. Not only are the cell side-walls flat, but so too are those that form the base of cells; base faces which are shared with cells on the opposite face of the comb. The mechanism by which bees build cells with flat sides has been the subject of speculation for centuries, but it has been conjectured by Kepler, Darwin as well as more recent researchers that bees build cylindrical cells that are transformed into flat sided prisms, without consensus as to the process by which this is achieved. By offering bees shaped wax stimuli and observing the comb that was built upon them, we have shown that under certain conditions walls will be curved and others where walls are reformed to be flat. A wall of a cell, be it a side-wall or a face of the base, with no other cell beyond it will be built as a convex curve whereas a wall with a cell to both sides will be formed flat. Furthermore, we show that these walls are plastic; walls that were initially built curved were re-shaped to be flat once a second adjacent cell had been built.

## Introduction

Honeycomb comprises recognisable hexagons, each with straight sides. Not only are the cell side-walls flat, but so too are those that form the base of cells which are shared with cell on the opposing face of the comb. The mechanism by which bees build comb, especially cells with flat sides, has been debated. It is our hypothesis that bees attempt to construct cells with a rounded, convex profiles and that the interaction between the attempt to form two such convex surfaces in contact with each other results in an equitable compromise – the flat face common to two cells. From this hypothesis I derived two predictions:

P1 - A wall with a cell to just one side will be curved, externally convex.

P2 - A wall with a cell to both sides will be straight; either formed straight or became straight.

These predictions were tested by measuring the curvature of isolated walls, those with a cell to just one side. Isolated walls either appeared spontaneously at the edge of comb construction, or as a result of offering the bees purposefully prepared stimuli where access to one side had been restricted. Following further construction of the comb, second stage measurements were made of shared cell walls, those with a cell to either side. Isolated walls, at stage 1 were found to be significantly more curved than shared walls and previously curved, isolated walls, were found at stage 2 to have become significantly straighter. These results support both predictions indicating that bees would build simple curved cells, but walls are flattened through interaction with builders forming adjacent cells. This supports the hypothesis that the facetted nature of cells and the architecture of honeycomb are emergent features arising from simple local actions.

Two noteworthy features of honeycomb are the isometric cells that form a regular pattern and the straight walls between each cell. It is the origin of the latter feature that was explored by four experiments i to iv described in this chapter. Cell walls, when first formed, have been observed to approximate to an arc of a circle and within older cells the sides appear to be straighter (Hepburn, Pirk, and Duangphakdee 2014:243). An attempt was made to explain this transformation (Pirk et al. 2004) but Pirk’s suggestion of thermoplasticity as the mechanism was rebutted (Bauer and Bienefeld 2012). Nonetheless the papers from Pirk et al. (2004) and Hepburn et al. (2014:243) contain the observations, now unexplained, that over the period of comb construction a cell wall becomes straighter. The purpose of the first stage of these experiments is to test whether early-stage cells are curved, and if so, whether this is a feature common to early-stage cells or just of walls with a cell to one side. Differing curvature in early-stage cell walls between those that are isolated and those that are shared would support my hypothesis and contribute to an explanation of cell formation. These experiments established the conditions for isolation of cell walls in four ways: cell location at the edge of comb, thick wax preventing access to cell side walls, physical barrier preventing access to the cell bases and thick wax preventing access to cell bases.

The method used by bees to manipulate the fabric of the cells is by small-scale manipulation, sculpting (Martin and Lindauer 1966), and in this fashion the bees can fabricate the walls of a cell from just one side. The purpose of these experiments is to demonstrate that single-sided construction is sufficient to form a curved cell, but not sufficient for the formation of facetted cells.

Hypothesis III, that construction workers will attempt to maximise the space within a cell, when applied to cells that share a wall, explains that the equilibrium between the attempt to form a convex shape by workers in adjacent cells gives rise to straight inter-cellular walls. A wall that, from the beginning, was formed straight would have been continually subject to the construction effort in each of the two cells; an influence that could have been simultaneous or alternated intermittently. Alternatively, a wall that had initially been built curved would, according to the hypothesis, subsequently be re-shaped, becoming flat during construction of the adjacent second cell. Re-formation of walls, changing from curved to flat, is the outcome expected by prediction P2 - A wall with a cell to both sides will be straight; either formed straight or became straight. This will be tested by a continuation of each experiment through to a second stage, measuring the result of continued construction on cell walls after the access restrictions had been lifted from each of the four test conditions.

For these experiments, we fashioned stimuli comprising wax forms designed to trigger the predicted behaviours. WeI then placed the stimuli in hives leaving the bees to build honeycomb upon them. Periodically, these samples were removed from the hives for inspection and photographed to record their current condition. The photographs were subsequently inspected to identify features that met the experiment specific criteria (set out below 0) and the curvature of those cell structures was measured. The walls that were the subject of these experiments included side walls, those between cells on a single face of the comb, and base walls, those between cells on opposing sides of the comb.

## Methods

### Hive handling and recording

#### Hives

Three British Nation hives housing locally reared honeybees, *Apis mellifera* were used in these experiments. The frames carrying the experimental stimuli were positioned at the edge of the brood zone.

#### Instigation of comb construction

Comb construction was encouraged by *ad libitum* feeding with 1:1 sucrose solution.

#### Wax

Wax used to form the various stimuli for these experiments had been recovered during previous seasons from hives within the same apiary was formed into sheets of either 0.5 mm-0.6 mm or 1.0 mm-1.2 mm thickness.

#### Preparation of stimuli

Stage 1 of these experiments tested prediction P1, that bees should attempt to build a cell shaped as a cylindrical tube and, where this construction is not constrained, the isolated cell walls (those with a cell to just one side) will be curved, externally convex. The four experiments (i to iv) each tested the prediction for walls that were isolated by different conditions. If the prediction is correct then, for all four conditions, the curvature of isolated walls will exceed that of shared walls more than expected by chance.

Stage 2 of these experiments tested prediction P2, that bees should attempt to build a cell shaped as a cylindrical tube and, where one such construction is conjoined to another, the common wall will be flat; a wall with a cell to both sides either will be formed straight or will become straightened. If the prediction is correct then, for all four conditions, previously isolated walls will have become curved significantly less compared with their previous state than expected by chance.

#### Preparation of stimuli for experiment i – side wall, progressive building

The first stage of this experiment causes the construction of isolated walls which subsequently, at stage 2, become common to two cells.

Where comb construction progresses normally, the cells at the periphery include walls that are at the edge of the comb beyond which there are no further cells. Toward the edges of naturally constructed comb, the cells, having been started only recently, are shallow and their height tapers to zero at the edge (Hepburn 1983). As a result of preliminary studies we noted that when such cells are still extremely shallow, there is little to distinguish between the bases and walls which would cause difficulties in measuring the shape of isolated walls, i.e. the walls of cells that had yet to have another cell built adjacent to it. Nonetheless, some cells that were close to the edge of comb were sufficiently distinct, typically being slightly less that one cell’s width from the edge and separated from the edge by an incomplete cell.

The stimulus used for this experiment was a flat piece of wax approximately 25mm by 40mm and 0.5mm thick, the surface of which was disrupted by the addition of additional pieces of wax or by indentations. These surface features had been found, during exploratory experiments, to promote the formation of comb onto the wax backplane. As cells were built onto the backplane, the comb commonly extended beyond the tab, providing some close to the edge to be sampled both when close to the edge, and subsequently after further cells had been built beyond the stage 1 sample walls (Figure 0-1). The curvature of peripheral walls when either isolated or shared, could be measured.

**Figure 0-1.**
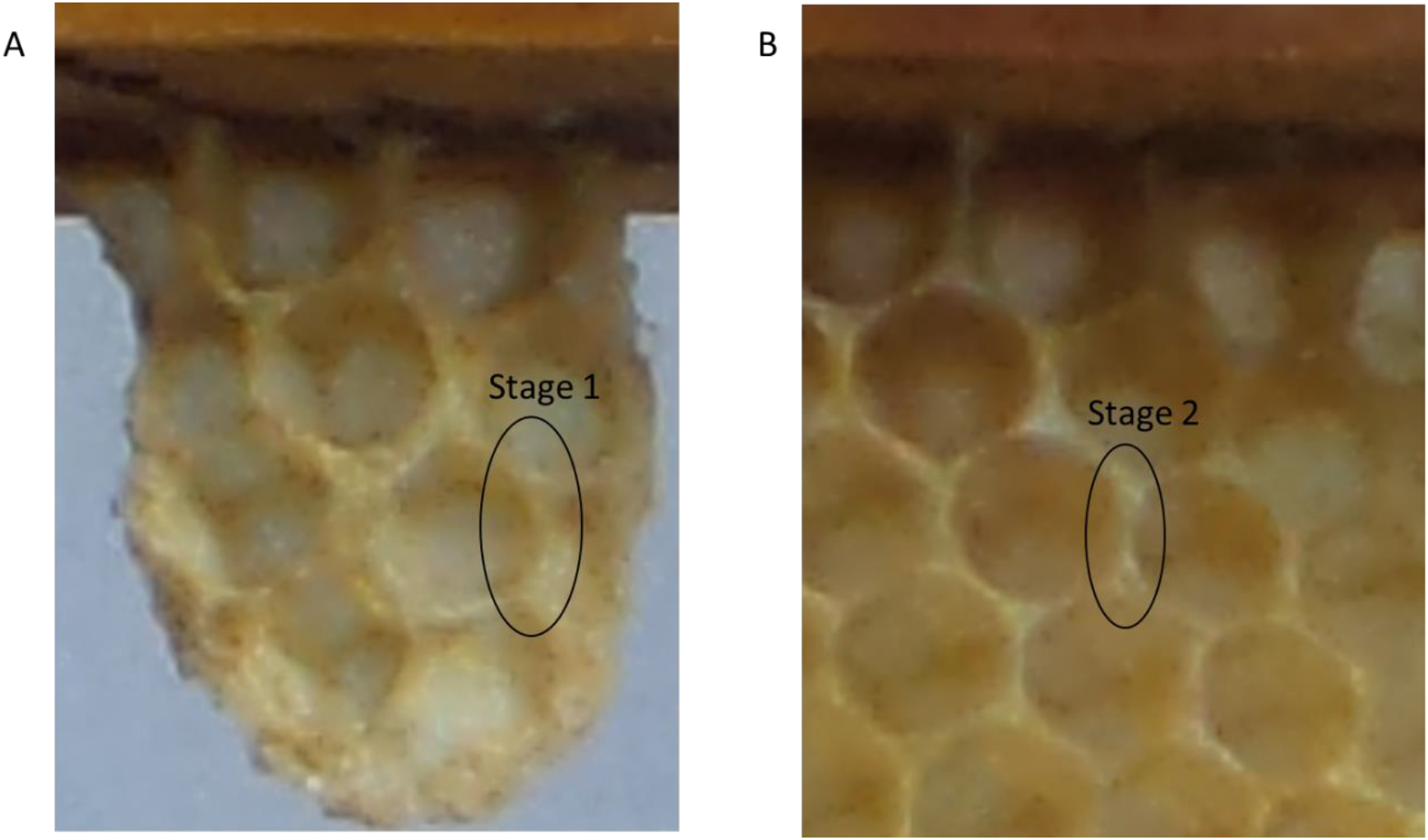
Illustration of experiment i. A: an example of a peripheral cell and a wall, marked “stage 1”, with a cell to just one side, at the earlier state at the edge of comb construction. B: a subsequently image shows the same cell, and wall, marked “stage 2”, after further cells had been built adjacent to it.

Stimuli prepared for this test were bonded to the upper bar of a test frame and placed within one of the hives. Test frames were inspected every three to four hours and photographed to record their current condition. The purpose of this experiment was to study cell walls as construction progressed, therefore these test pieces were returned to the hive for further construction for as long as there were distinct edges to the honeycomb.

The second stage of this experiment was to test prediction P2 using cells that previously had at least one wall that was not shared with another cell and where a second cell had subsequently been built to share that wall.

The test pieces, previously used for stage 1 of this experiment, were replaced within each hive and construction was allowed to continue. Test frames were inspected every three to four hours and photographed to record their current condition, thus a wall that had been measured when isolated could be re-measured at the second stage once an additional cell had been built to the other side of the wall. Measurements were taken only for walls where the additional cell had been built to a depth of at least 5mm.

#### Preparation of stimuli for experiment ii - side wall, thick wax

This experiment involved cells to be built where at least one wall of the cell would be formed against a thick section of wax and which subsequently, at stage 2, had become common to two cells.

The test pieces were fabricated using a backplane, approximately 25mm by 40mm and 0.5mm thick, onto the face of which seed strips were added. The seed strips were 2mm-3mm in height and 1-1.2mm width and had been prepared by cutting strips from a flat sheet of wax formed by the dipping method (see 3.3.1.3), doing so 6 times in this case. The seed strips were welded onto the wax backplane and small depressions, of 4mm diameter, were pressed into the backplane close to the strips as further encouragement for the construction of cells close to the strips.

Stimuli prepared for this test were bonded to the upper bar of a test frame and placed within one of the hives. Test frames were inspected every three to four hours and photographed to record their current condition. Test pieces were returned to the hive for further construction while the subject cells, those against the barrier, were shallower than 5mm.

The purpose of the second stage of this experiment was to test prediction P2 applied to cells that had at least one wall that had been formed by a thick section of wax and where the barrier was subsequently thinned, and a second cell then shared that wall (Figure 0-2).

**Figure 0-2.**
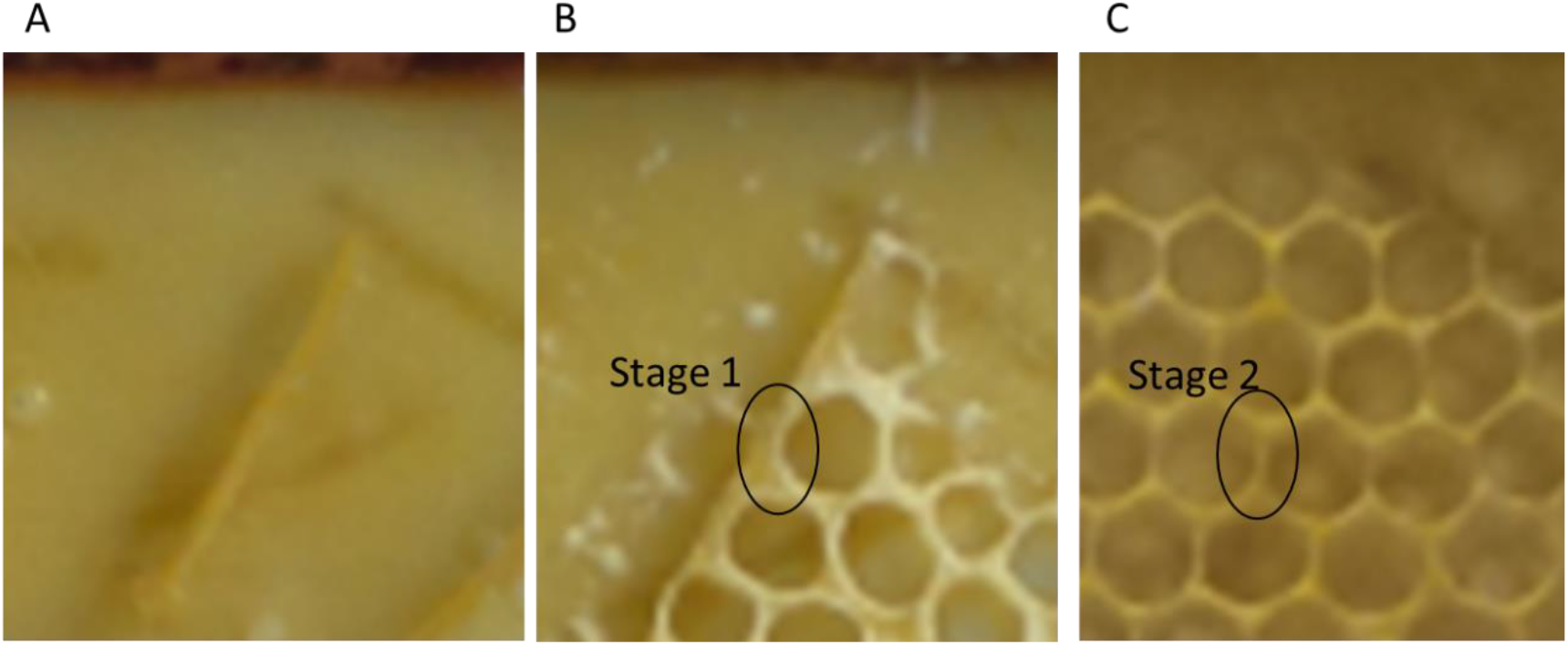
Illustration of experiment ii. A: initial stimulus formed of a wax strip welded to the tab, or backplane. B: at the first stage the circled cell is an example of one with a wall built against, and partially into, the thick barrier. At this stage, the wall had been constructed from just one side so is curved. C: a subsequently image shows the same cell and wall after further cells had been built and the barrier had been eroded. At this later stage, the wall had become subject to construction on both sides and had become straight.

The test pieces were replaced within each hive and construction was allowed to continue. Test frames were inspected every three to four hours and photographed to record their current condition, thus a wall that had been measured when isolated could be re-measured at the second stage once the thick wall had been eroded, allowing an additional cell to share the wall. A wall would be suitable for a second measurement provided that the cells to both sides had reached a depth of at least 5mm.

#### Preparation of stimuli for experiment iii - base, access denied

This experiment was performed to test prediction P1 applied to the bases of cells where the construction workers had access to only one face of the base walls. The second stage of this experiment allowed access to both faces of the cell bases to test prediction P2.

The configuration during stage 1 allowed comb construction to proceed on just one face of the, normally double-sided, comb by preventing access to the opposite face of the comb. The test pieces were fabricated using a backplane, approximately 75mm by 40mm and 0.5mm thick, into one face of which small depressions, of 4mm diameter, were pressed. Each sheet of wax was bonded onto the top bar of a frame, using molten wax. A similar sized (75mm x 40mm) piece of acrylic sheet was bonded to the frame, also using molten wax. The acrylic sheet barrier was positioned parallel to the wax sheet, separated by 3mm-4mm. Small wax columns of approximately 3mm diameter, formed from drops of molten wax, were placed at the two unsupported corners thus securing the wax sheet to the barrier. This was necessary as, otherwise, the construction workers had proven themselves quite capable of enlarging the gap to gain access and so form comb on both sides of the wax.

Stimuli prepared for this test were bonded to the upper bar of a test frame and placed within one of the hives. Test frames were inspected every three to four hours and photographed to record their current condition. Test pieces were returned to the hive for further construction while the subject cells, those opposite the barrier, were shallower than 5mm.

Stage 2 of this experiment was performed to test prediction P2 applied to cells where access to the base had been blocked and where the barrier was subsequently removed allowing a second cell to be constructed adjacent to the base.

The test pieces, used for stage 1, were adjusted to remove the barrier from half of the backplane and enable access there to both sides of wax where cells had hitherto been built to just one side. For the other part of the wax the barrier continued to prevent access. After the barrier had been adjusted, the frames were replaced within each hive and construction was allowed to continue.

Test frames were inspected every three to four hours and photographed to record their current condition, thus base walls that had been measured when access had been blocked could be re-measured at the second stage, once access had been possible, allowing cell on the other face of the backplane to share the base. Bases would be suitable for a second measurement provided that the cells to both sides had reached a depth of at least 5mm.

#### Preparation of stimuli for experiment iv - base, thick wax

This experiment was performed to test prediction P1, that isolated cell walls (those with a cell to just one side) will be curved, applied to the bases of cells where the influence of construction workers in cells on opposite sides of the comb were isolated by a thick layer of wax. The second stage of this experiment allowed continued construction, eroding the wax and so subjecting the bases to dual-sided influence to test prediction P2, that the double-sided influence will cause walls to become straight.

The first stage created conditions where the construction workers could influence the cell bases from inside the cells only. The configuration allowed comb construction to proceed independently on both faces of backplane. The wax that formed the backplane was thick enough to prevent construction work on a base from within a cell on one face of the comb influencing that being done on the base of a cell on the other face. Exploratory tests had shown that the thick wax provided isolation of one base from another for long enough for the bees to form hexagonal, flat sided cells of approximately 5mm depth. When allowed to continue to a cell depth of approximately 8mm, the wax at the base had been eroded sufficiently to allow the two sides to interact. Therefore, cells sampled for the first stage of this experiment were those of between 4-6mm depth and those greater than 8mm depth were chosen for the second stage.

The test pieces were fabricated using a backplane, approximately 75mm by 40mm and 1mm-1.2mm thick which had been prepared by a flat mould into molten wax, dipping the mould six times in this case to achieve the required thickness. Both faces of the backplane were treated with shallow depressions, of 4mm diameter. Each sheet of wax was bonded onto the top bar of a frame, using molten wax.

Stimuli prepared for this test were bonded to the upper bar of a test frame and placed within one of the hives. Test frames were inspected every three to four hours and photographed to record their current condition. Test pieces were returned to the hive for further construction while the subject cells, those above the barrier, were shallower than 5mm but less than 8mm.

The purpose of the second stage of this experiment was to test prediction that dual-sided access leads to straight walls (P2) applied to cells where access to the base had been limited by a thick wax substrate and where the wax had eventually been eroded allowing the base walls to be subject to influence from both sides.

Test pieces, previously used for stage 1 causing curved walls to be built, were reused and the frames were replaced within each hive and construction was allowed to continue. Test frames were inspected every three to four hours and photographed to record their current condition. Bases would be suitable for a second measurement provided that the cells to both sides had reached a depth of at least 8mm.

#### Stimulus handling and construction time

The frames carrying experiment stimuli were placed within hives for periods of three to four hours, repeated until construction had proceeded sufficiently.

### Measurement methods

#### Recording and photography

Progress of construction was recorded by photographing the test frames between sessions within the hive.

#### Photographic record analysis

We wrote custom software, FormImageCompare, to align and compare pairs of images. This software included a number of options that provided the means to mark and measure features of the stimuli and comb, thus yielding the data to produce the experimental results.

### Measurement methods

Thesis predictions P1 - a wall with a cell to just one side will be curved - and P2 - A wall with a cell to both sides will be straight - both concern the curvature of cell walls, either side walls or bases. The profile of each side wall was measured from the photographs recording the face of the comb. Base curvature measurements were obtained with the aid of a profile probe.

#### Measurement of side wall curvature

A cell’s side walls were measured from the photographic records using the coordinates of points marked on photographs taken of each face of the subject comb. The photographs recorded each side of the test pieces before placement in the hives, and at each stage of inspection. The frame comparison tool, FormImageCompare, includes a feature to measure ‘curves’. This feature allows the user to mark, by mouse click, two points at the ends of a wall section and the wall midpoint – see Figure 0-3. This is done initially when viewing the first stage photograph, then a second time for the second stage image. The zoom feature of the software allowed the images to be expanded, assisting the placement of the measurement points.

**Figure 0-3.**
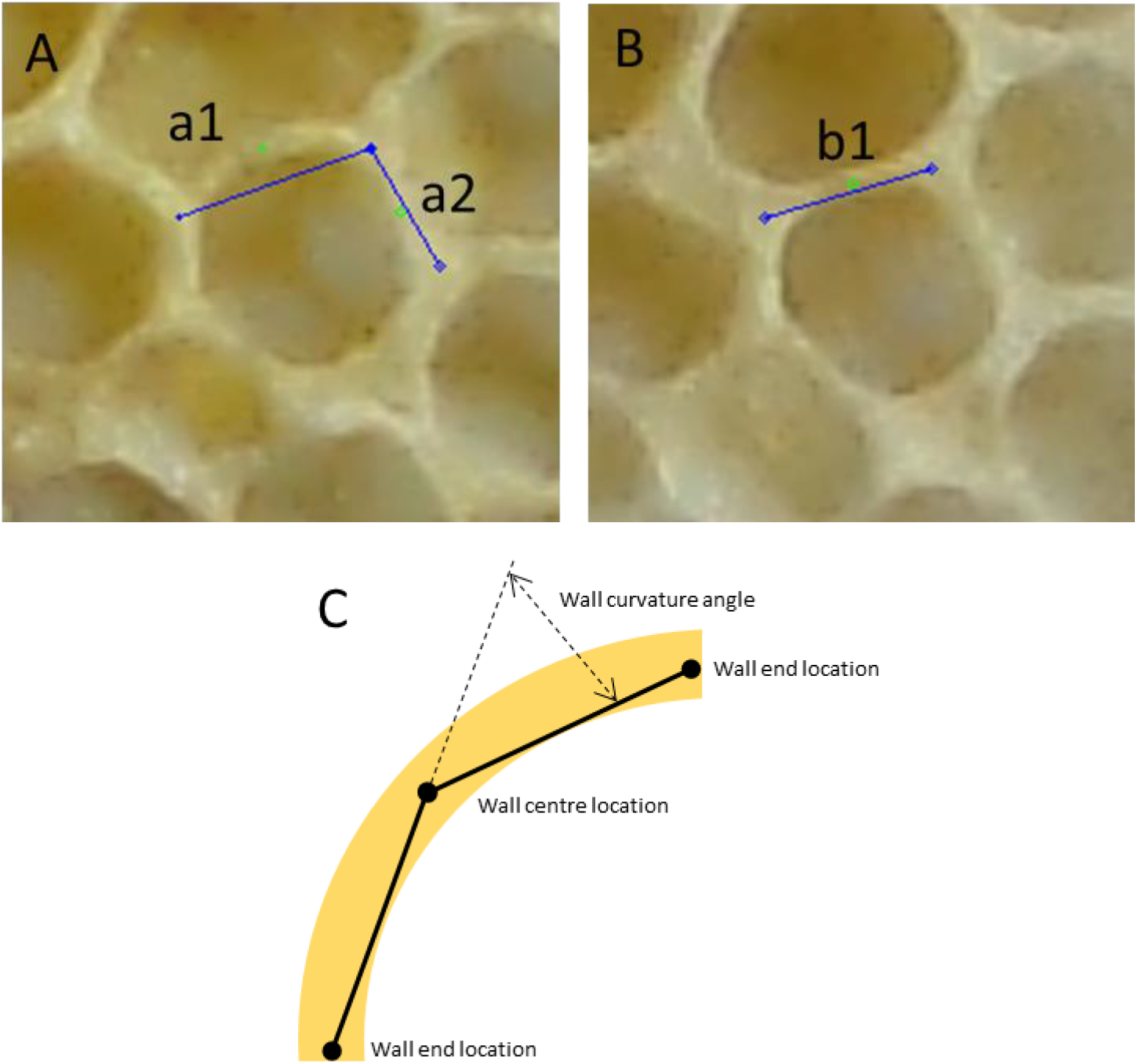
Placement of marks used to measure the curvature of side walls.A: the form of a cell at its early stage. The upper wall, a1, had a cell to just one side and was measured as an isolated wall at stage-1 while the wall a2, common to two cells, was used as a control sample. B: the same section of comb following further construction and the wall, b1, was measured as a sample of a shared wall at stage 2. The cell wall measurements were computed using the coordinates of the manually placed marks. Blue diamonds mark the corners of walls and blue lines join the corners illustrating where a perfectly straight wall between those corners would run. Green circles represent the mid-point of each wall. C: diagram showing example wall marks. The metric of curvature used for anaylis the difference in angle between the two lines; from the centre to each end.

The quantity used to describe the curvature of each cell wall was the angular difference between the line from the centre of the wall to a point at each end of the measured structure.

#### Measurement of base wall curvature

The curvature of a cell’s base wall is not easily visible without cutting through the comb to expose the base profile. Sectioning the cells was not chosen due to the disturbance likely to be caused to the bases reducing the accuracy of results obtained that way. Also, testing of the prediction that further construction work would straighten a curved base could only be achieved if a non-destructive method were to be used for the first measurement.

A profile gauge was constructed comprising seven depth probes bonded into a linear array so that the whole width was sufficiently narrow to fit within a worker cell without distorting the cell walls.

Each probe was made using an outer barrel with a sliding inner wire. The barrel was a syringe needle from which the plastic mounting collar was removed. The needles used were 23-gauge with outer diameter of 0.64 mm and bore diameter of 0.34 mm, The inner wire was a piece of 10-gauge guitar string, diameter 0.010 inch (0.245 mm). The seven needles were bonded into the array with epoxy resin (Araldite) forming a contour probe with an overall width of with 4.9 mm. The probe wires protrude from the core of the guide tubes and therefore the span of the probes was 4.6mm.

The assembly included a gauge comprising lines separated by 1mm printed on paper mounted to the rear of the top end of the probe.

Base profile measurements were taken by first extending the probe wires so the top end of each wire was approximately flush with the top of the probe barrels. The probe assembly was pressed into the cell, following the line of the cell side walls to align the device to the cell. When possible, the probe state was recorded by photographing the top part of the gauge while in-situ. In some cases, the angle of comb, alignment or width of the cell prevented the probe from resting within the cell stably enough to permit photography. On these occasions the probe was withdrawn, and the bottom part photographed alongside a 10mm gauge.

A custom software tool, FormDepth, was used to extract numeric values from the photographic records. We wrote FormDepth in C++ using Microsoft Visual Studio Community 2019:Version 16.7.2, Visual C++ 2019, drawing on support from the library OpenCV:Version 3.3. This tool is available at https://github.com/VinceGalloQMUL/honeycombThesisRepo.

The tool allows the photograph to be viewed at various degrees of magnification which ease the task of placing a marker on the tip of each probe wire. Two other marks, placed at each end of a 10mm reference object, provide the software with a calibration distance. The distance reference was either the printed label built into the gauge, used for measurements using the top of the pins, or the jaws of vernier callipers opened to 10mm placed within the field of view for measurements taken from the lower ends.

The software used the distance reference locations to calculate the image scale in terms of pixels per millimetre. The scale, computed separately for each image, was used by the software to convert the locations of the marks at each probe into a height for each probe pin. Absolute heights were not used, rather the left-hand end probe was used as a reference point with the distance above or below that datum being recorded for the other probes.

For measurements using the top of the probe pins it was necessary to first calibrate the lengths of each probe. The probe was used several times to ‘measure’ a flat surface, and the relative heights used to compute a mean value of the offset to be used for each probe. The resulting calibration values were then used to adjust measurements based on the top of the probe pins.

Measurements of the bases using the multi-point depth gauge were used to determine whether the degree to which bases were curved, or faceted.

The measurements obtained from the photograph yielded the depth at 7 points (d_0_-d_6_) across the cell relative to pin 0 (d_0_ = 0). The depths were first adjusted to level the profile and thus correct for inclination of ether the cell base or the insertion of the probe. Levelling was done by transforming the measurements into values relative to a straight line between d_0_ and d_6_. Two values were then computed from the levelled profile to derive a metric of being curved or facetted.

A curved base was assumed to be a section of a sphere and the profile gauge to have sampled the spherical profile along a linear cross section, hence the measured profile would for an arc of a circle. The measure of curvature ‘S’ was calculated as the sum of the squared deviation of each measured height from the corresponding point on a circular arc through points d_0,_ d_3_ and d_6_.

The measure of deviation from a faceted profile ‘F’ was calculated as the sum of squared deviation from two regression lines fitted to the points from d_0_ to the lowest point and from that point to d_6_.

The metric of base curvature ‘C’ was the degree to which the profile matched that of a curved base compared with that to a facetted base.

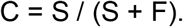

#### Base curvature measurement error estimation

Errors in the measurement of base profile could arise in several ways including physical aspects of the gauge, handling of the gauge and errors by the user when marking locations using the software. An estimate of the total errors that might arise from this measurement was obtained by repeated measurement of a single cell. The cell selected for repeated measurement was chosen from those formed on the thick backplane (section 0) as its substantial base would resist repeated probing.

### Experiment specific measurements

#### Experiment i - side wall, progressive building

Experiment 4i tested predictions P1 and P2 in the circumstances where comb had been spreading, resulting in cells at the edge of that comb. For this experiment, isolated walls found at the edge of comb were selected, but only those for which a subsequent photographic record would permit their inclusion in the second stage measurement.

The control population comprised, for each sample cell, a second contemporaneous wall for which there was already an adjacent cell. The control walls were selected by first considering a wall in the same cell as, but opposite to, the experimental isolated wall. If the candidate control wall was too indistinct to be measured, then the alternative control wall was selected. The alternative wall would be found in the next cell, traversing the comb via the initial candidate control wall. The control wall would then be chosen as that opposite the initial candidate.

Stage 2 of this experiment re-measured the curvature of the isolated walls that had been sampled in stage 1, after subsequent construction of additional cells, when the walls had become shared. The additional cell, for this purpose, would be a complete convex enclosure; corners comprising only obtuse angles

#### Experiment ii - side wall, thick wax

Cell walls sampled for stage 1 of this experiment were those that were insulated from the influence resulting from the construction of an adjacent cell by being built against a thick wax barrier. For this experiment, cell walls built against the wax barrier were suitable when construction had progressed to the point where an identifiable cell had been formed – enclosing at least 270°, but where no cell had yet been built on the other side of the seed strip. A further condition for selection was for a subsequent photographic record suitable for inclusion of the wall in the sample set for the second stage measurement. Hence, the sampled walls were those that abutted the stimulus barrier and later, after further construction, became shared by a cell built to the other side of the barrier.

The control population was sampled using the same method as described for experiment i.

Stage 2 of this experiment re-measured the curvature of isolated walls that had been sampled in stage 1, after subsequent construction of additional cells, when the walls had become thin and shared.

#### Experiment 4iii - base, access denied

Cell bases sampled for this experiment were from an area where access to the reverse side of the stimulus had been prevented by a physical barrier. The measurements were made using the profile gauge as described in 0.The subject bases used for the initial measurements (subsequently referred to as “Stage 1”) were those of cells with wall heights of 4mm to 6mm. The natural form of honeycomb tends to be thicker in the middle, tapering to the edges, which meant that most candidate cells were those built around the middle of the stimulus backplane.

The control population (subsequently referred to as “Natural”) comprised cells of 4mm to 6mm depth, that formed part of natural comb build onto the same frame, but away from the barrier.

The second stage tested prediction P4.2 in the circumstances where access to both faces was possible, having previously been prevented. Further measurements of the cell bases were made of cells where construction had proceeded to the point where cells on uncovered sides had achieved a depth of between 5mm and 8mm. Measurements were taken using the gauge to measure the profile of cell bases. Two sets of cells were sampled; cells within the newly uncovered area (described as “Stage 2 – uncovered”), and cells within the area where the barrier had been retained (described as “Stage 2 – covered”).

#### Experiment 4iv - base, thick wax

Cell bases sampled for this experiment were from an area where comb had been built onto the thick wax backplane. The measurements were made using the profile gauge as described above.

The subject bases used for the initial measurements (subsequently referred to as “Stage 1”) were those of cells with wall heights of 4mm to 6mm. The natural form of honeycomb tends to be thicker in the middle, tapering to the edges, which meant that most candidate cells were those built towards the middle of the stimulus backplane.

The control population was that used as the control population for experiment iii subsequently referred to as “Natural”.

Measurements of the cell bases (described as “Stage 2) were made of cells where construction had proceeded so that cells been built to a depth of between at least 8mm.

### Analysis

Experiment i - side wall, progressive building

and

Experiment ii - side wall, thick wax

The curvature of isolated cell walls in the sample measured at stage 1 had a distribution which appears normal, but the distribution of values for the control samples and those measured at stage 2 were not normally distributed. Therefore a ranked test was used for the comparison, specifically, the R function wilcox.test().

At stage, 1 comparing the samples with the control set, a two-sample test was performed. When comparing the stage-2 measurements with those taken at stage 1 for the same walls, a paired test was used.

Equivalence was tested between the measurements obtained from cell sides for the three situations when access was possible on both sides (4i stage 2, 4ii stage 2 and control). This was done by performing a TOST comparison on each of the three possible pairings. The smallest effect size of interest (SESOI) used for these was 0.5 of the standardised mean difference (Cohen’s D).

Experiment iii - base, access denied

and

Experiment iv - base, thick wax

For these experiments, comparison of the stage 1 samples with the control set and comparison between stage 2 values and the previous, stage 1 values a Wilcoxon two-sample test was performed.

Equivalence was tested the measurements obtained from bases for the three situations when access was possible on both sides (4iii stage 2, 4iv stage 2 and control). This was done by performing a TOST comparison on each of the three possible pairings. The smallest effect size of interest (SESOI) used for these was the estimated measurement error of 0.105 (Base curvature measurement error estimation).

## Results

### Experiment 4i – cell side wall curvature was influenced by isolation or being shared during progressive comb building

We measured 62 isolated walls found on comb from 11 frames and a further 62 non-isolated walls as a control sample. Subsequently, at stage 2, I re-measured the previously isolated walls.

The curvature of the isolated walls was 0.219 radians ±0.076 (mean ± standard deviation, throughout), which was significantly greater than the curvature of non-isolated walls (0.070 radians ±0.053, W_61_ = 3625, P<0.00001; Figure 0-7 A & B). This demonstrates that cell walls curve where constraint imposed by an adjacent cell is absent, as predicted when one assumes that bees attempt to construct cells with a rounded, convex profiles.

**Figure 0-4.**
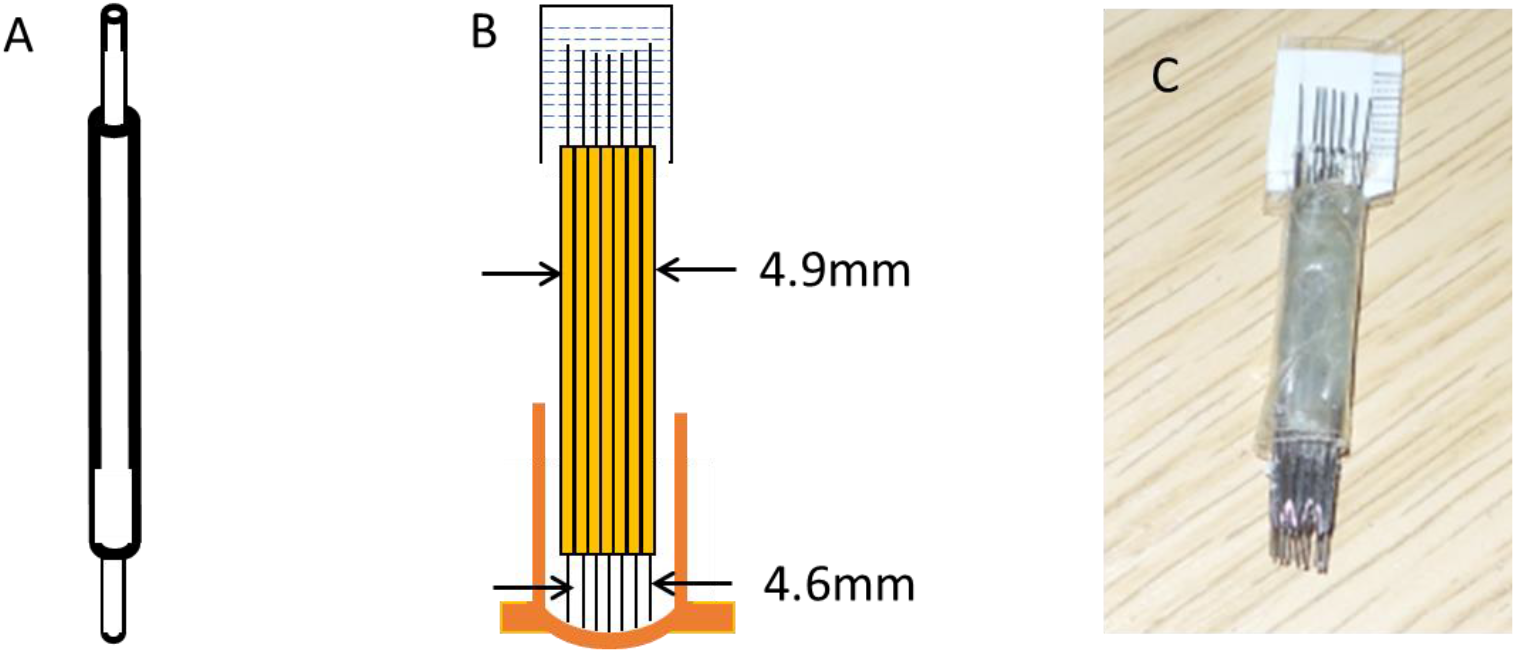
Depth gauge construction. A: diagram of a single hypodermic needle with wire insert. B: diagram of seven probe barrels glued together to form a profile gauge. C: photograph of the probe used for the experiment.

**Figure 0-5.**
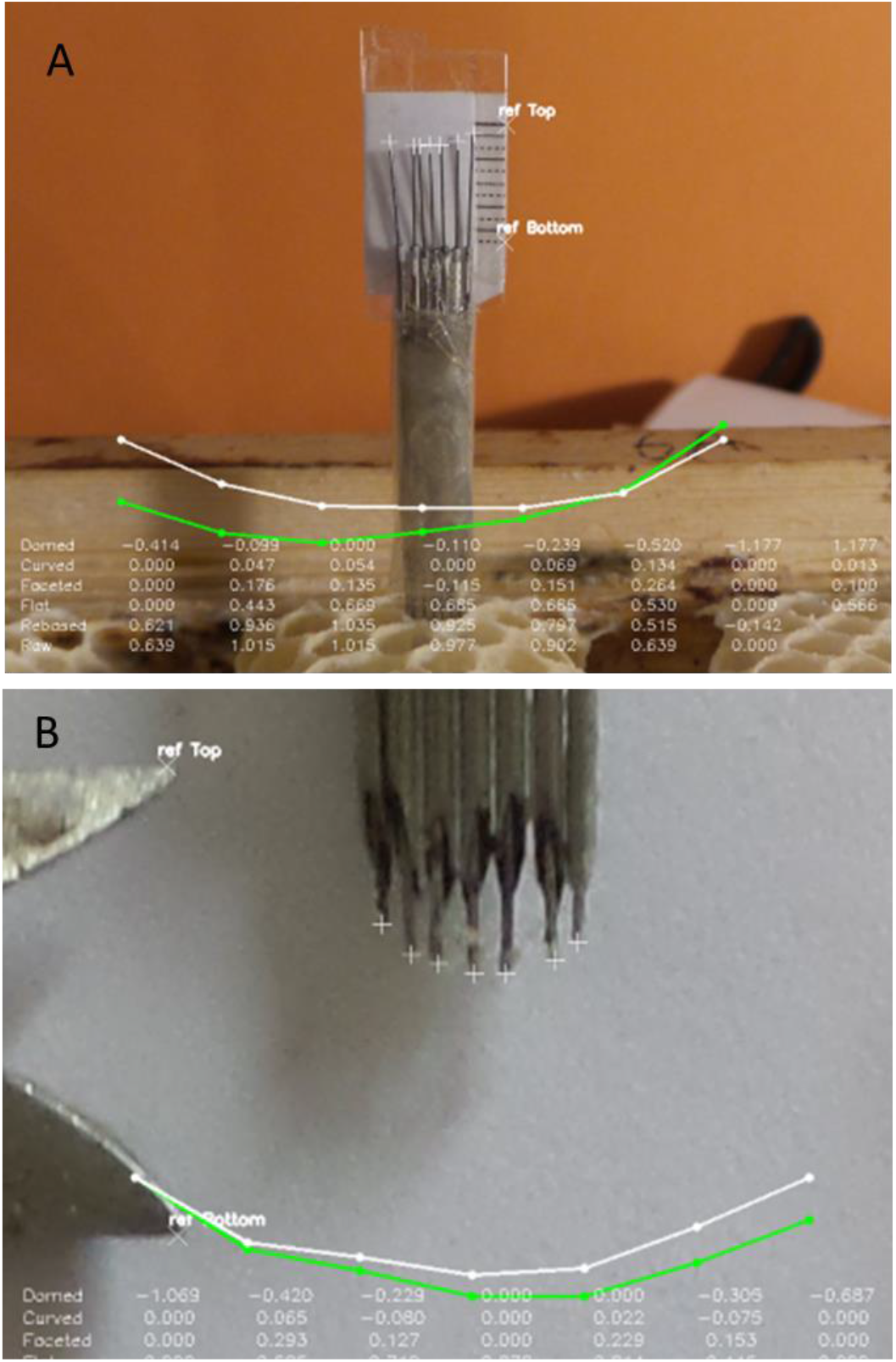
Depth gauge use and processing. A: photographic record with probe in situ. B: record of profile having been removed from the cell. In both images the gauge top and bottom marks were place by the user at either end of a 10mm reference. The ends of the probe wires were marked, manually, and the software used those locations, together with the gauge marks, to compute the relative heights of the seven profile points. The software provided feedback by drawing curves formed from seven points and interconnections. The green lines and points show the raw measurements while the white lines and points show the curve having levelled the two end points.

**Figure 0-6.**
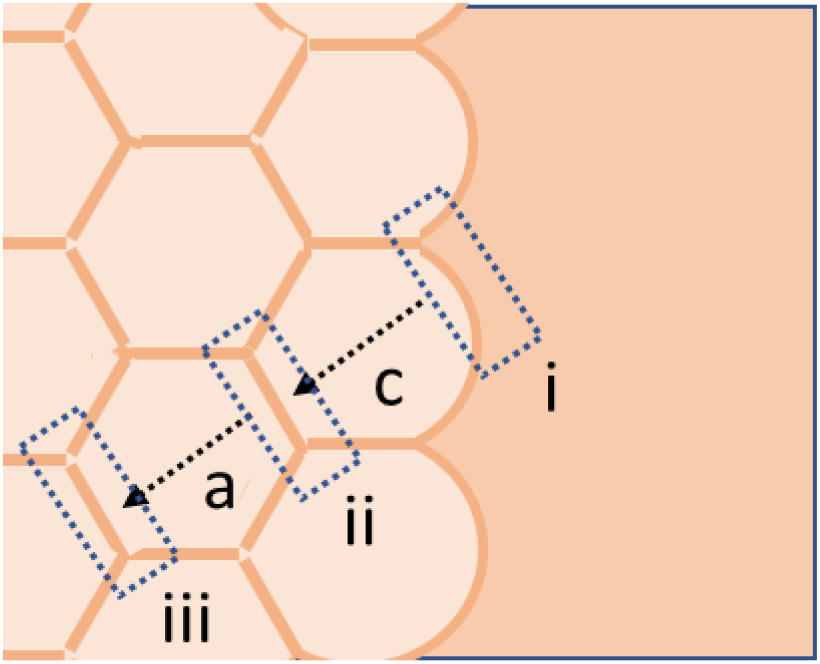
Isolated side wall and selection of control sample. The wall (i) on the outside of the peripheral cell (c) may be selected for inclusion in the set of sample isolated walls. In this case the first choice for a control sample would be the wall (ii) of cell (c) opposite wall i. The alternative, used when wall ii was indistinct, would be the wall (iii) on the opposing side of the adjacent cell (a).

**Figure 0-7.**
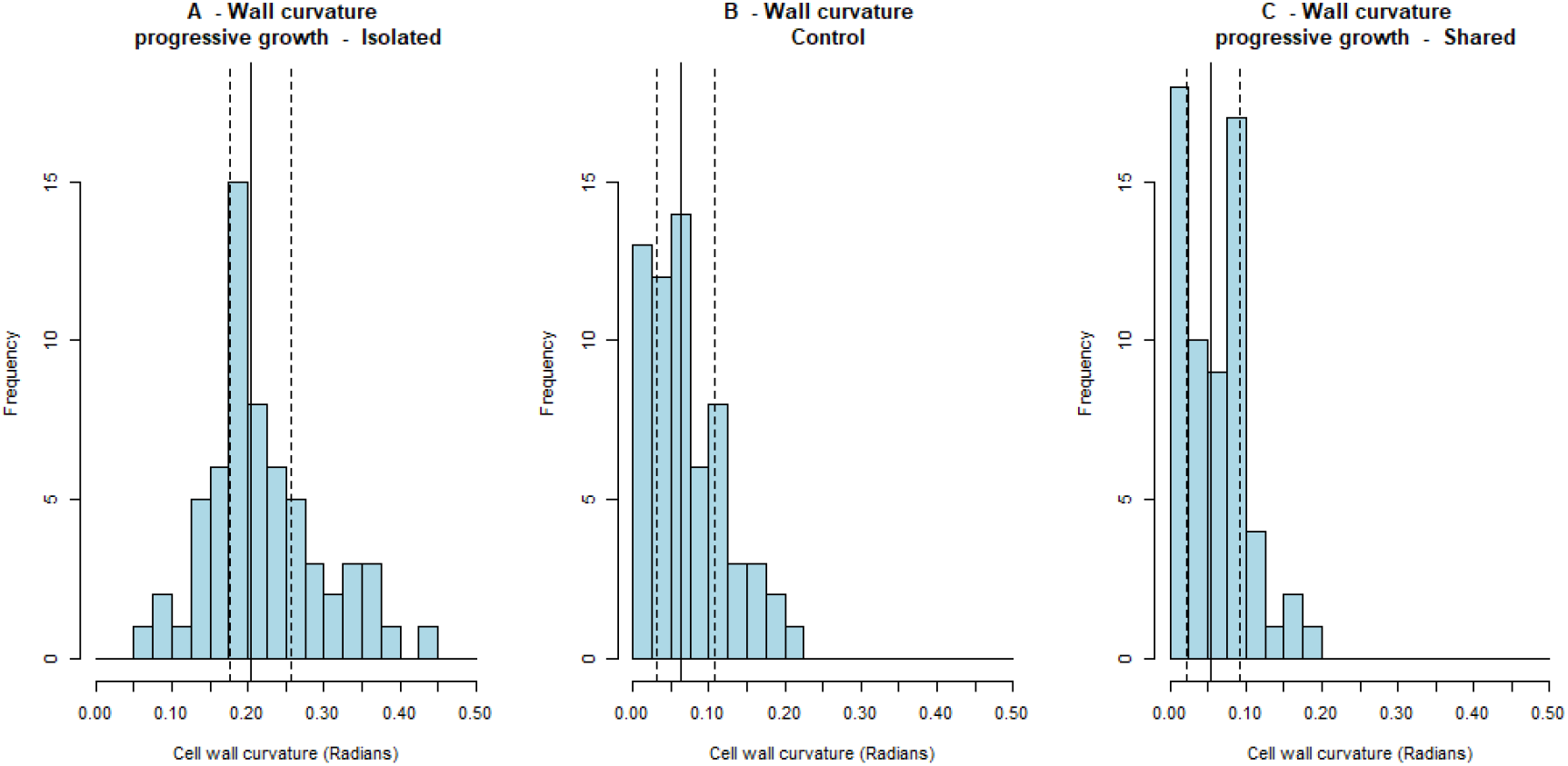
Curvature of stage 1 cell walls. A: the curvature of cells walls at the edge of a piece of comb, with a cell to just one side. Solid vertical line shows the median, dashed lines the interquartile range. N=62. B: curvature of cell walls with a cell to both sides. N=62. C: curvature of cell walls, previously with a cell to just one side, at stage 2 with a cell to both sides. N=62. Walls with cells to both sides are straighter than those with a cell on just one side.

Walls, initially isolated on the edge of developing comb (Figure 0-7 A) had a mean curvature of 0.219 radians ±0.076 across 62 samples. When re-measured, for this experiment, the curvature had reduced significantly (0.061 radians ±0.043, V_61_ = 1941, P <0.00001; Figure 0-7 C). This demonstrates that cell walls are straightened in the presence of an adjacent cell (Figure 0-8), as predicted (P2) - A wall with a cell to both sides will be straight; either formed straight or became straight - derived from the assumed interaction between the bees attempt to construct both adjacent cells with a rounded, convex profiles.

**Figure 0-8.**
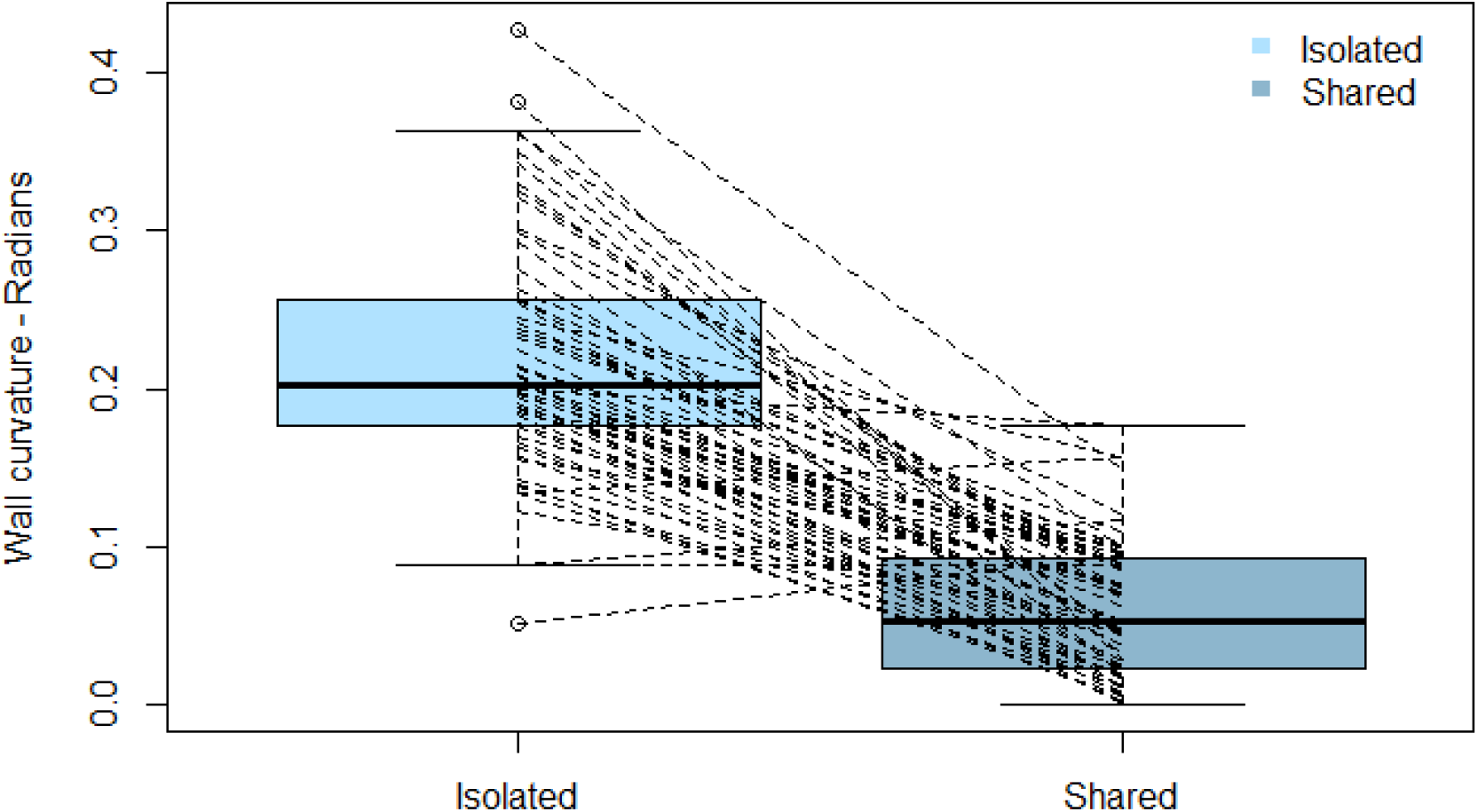
Cell wall curvature when isolated and when shared and the changes for each wall. Isolated walls, when there was a cell to just one side, compared with subsequent measurements taken when the walls each had a cell on both sides. The connecting lines from left to right on the chart show the individual transition for each wall.

### Experiment ii – curvature of cell side wall against a thick wax barrier was influenced by isolation or being shared

We measured 43 instances of an isolated wall abutting thick wax barrier and a further 43 non-isolated walls as a control sample.

The curvature of the isolated walls was 0.259 radians ±0.060, which was significantly greater than the curvature of non-isolated walls (0.129 radians ±0.061, W_42_ = 1723, P <0.00001;Figure 0-9 A & B). This demonstrates that cell walls curve where there is no constraint imposed by an adjacent cell, as predicted when one assumes that bees attempt to construct cells with a rounded, convex profiles.

**Figure 0-9.**
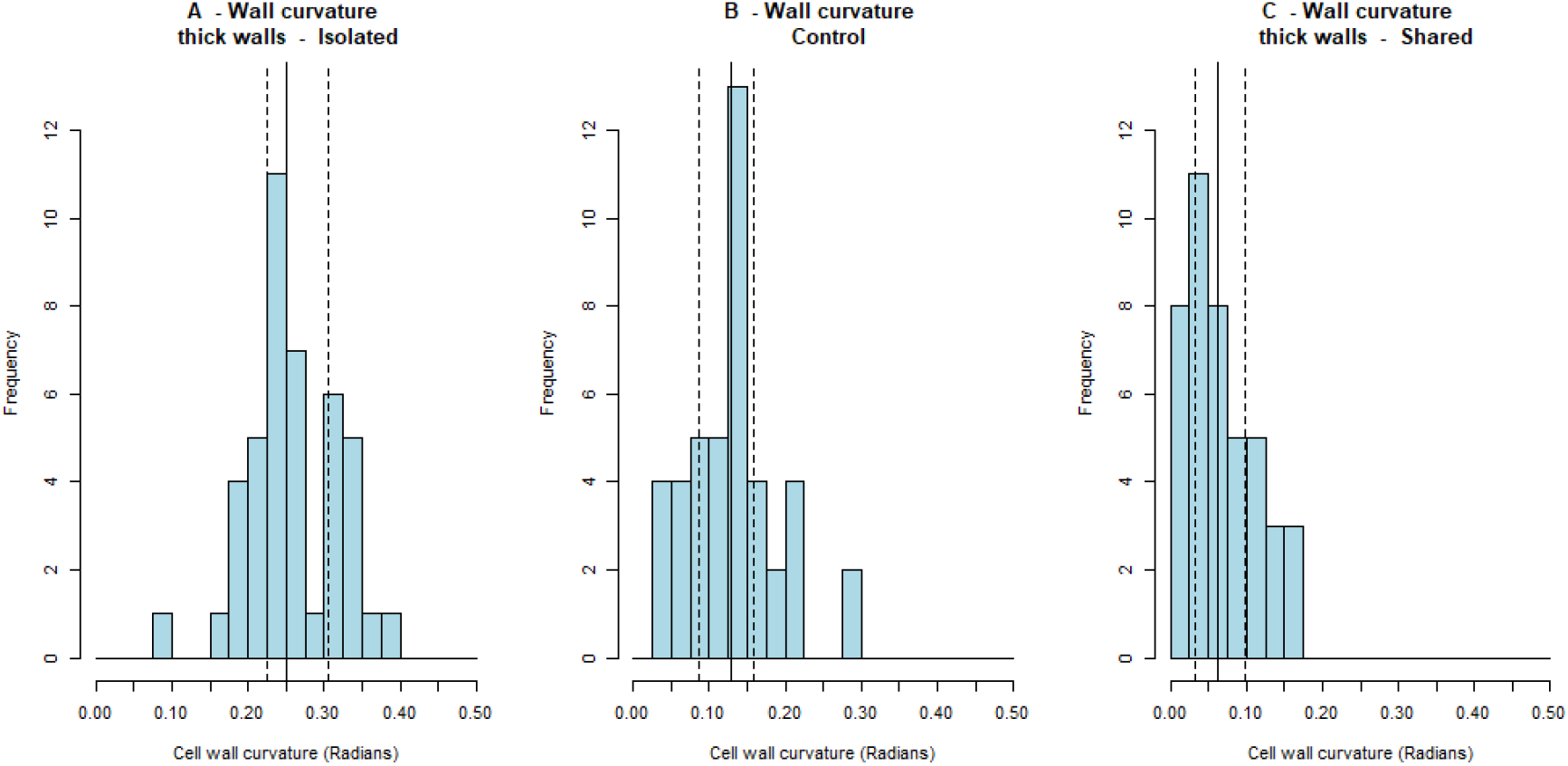
Curvature of stage 1 cell walls. A: the curvature of cells walls against a thick barrier. Solid vertical line shows the median, dashed lines the interquartile range. N=43. B: curvature of cell walls with a cell to both sides. N=43. C: curvature of cell walls at stage 2, once the thick barrier had been eroded, with a cell to both sides. N=43. Walls with cells to both sides are straighter than those with a cell on just one side.

Walls, initially part of a thick wax barrier, had a mean curvature of 0.259 radians ±0.060 across 43 samples. When re-measured, for this experiment, the curvature had reduced significantly (0.068 radians ±0.045, V_42_ = 943, P <0.00001; Figure 0-9 A & C). This demonstrates that cell walls are straightened in the presence of an adjacent cell (Figure 0-10), as predicted (P2) - A wall with a cell to both sides will be straight; either formed straight or became straight - derived from the assumed interaction between the bees attempt to construct both adjacent cells with a rounded, convex profiles.

**Figure 0-10.**
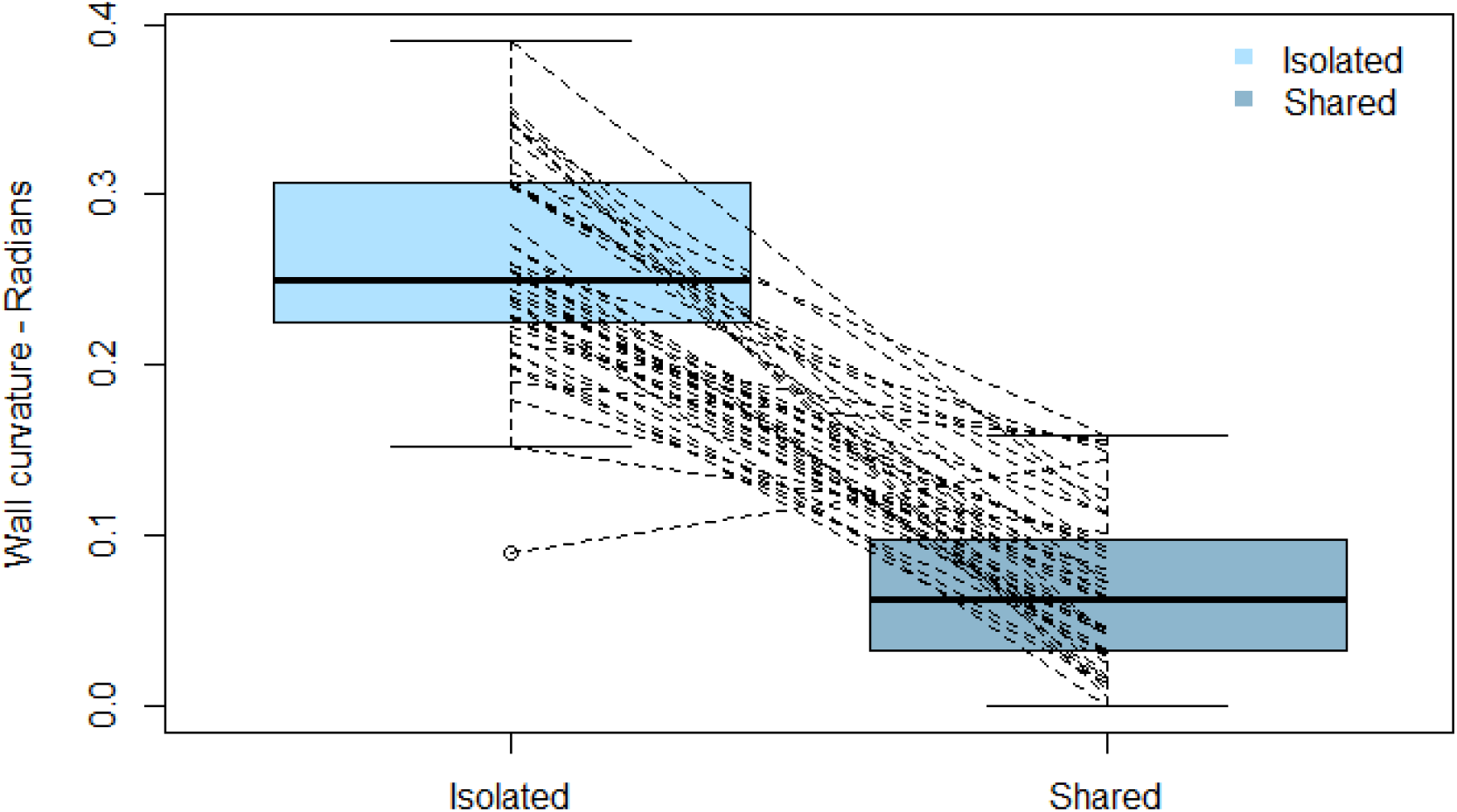
Cell wall curvature when isolated and when shared and the changes for each wall. Isolated walls were those against a thick wax barrier, compared with shared walls that were measured subsequently when the barrier had been eroded such that the walls each had a cell on both sides. The connecting lines from left to right on the chart show the individual transition for each wall.

### Experiment iii - cell base wall curvature was influenced by isolation or being shared

We measured 99 instances of cell bases accessible from only one side, access to the other side having been prevented by a physical barrier. A further 43 bases found within natural comb as a control sample. Later, at stage 2, We measured the bases of 31 cells that had remained covered and the bases of 50 cells where the protective cover had been removed.

The curve factor for the isolated bases was 0.74 ±0.18, which was significantly greater than the factor for natural bases (0.30 ±0.14, W_101_ = 4032, P <0.00001;Figure 0-12 A & B). This demonstrates that cell bases curve where there is no constraint imposed by an adjacent cell, as predicted when one assumes that bees attempt to construct cells with a rounded, convex profiles.

**Figure 0-11.**
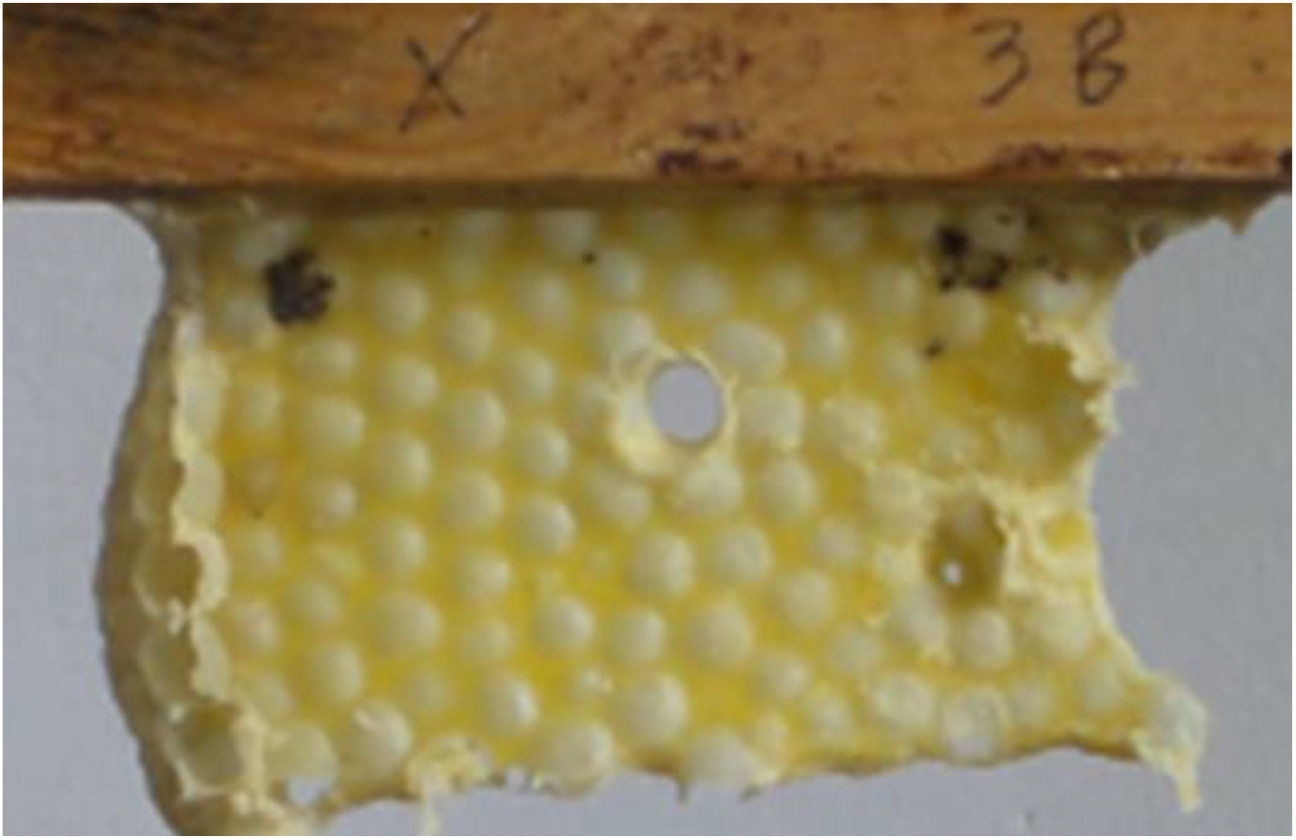
Underside of a sample where cells had been built onto backplane showing the face that had been protected by a barrier, since removed. The surface of the wax has been disturbed by the domed bases to cells built on the opposite face.

**Figure 0-12.**
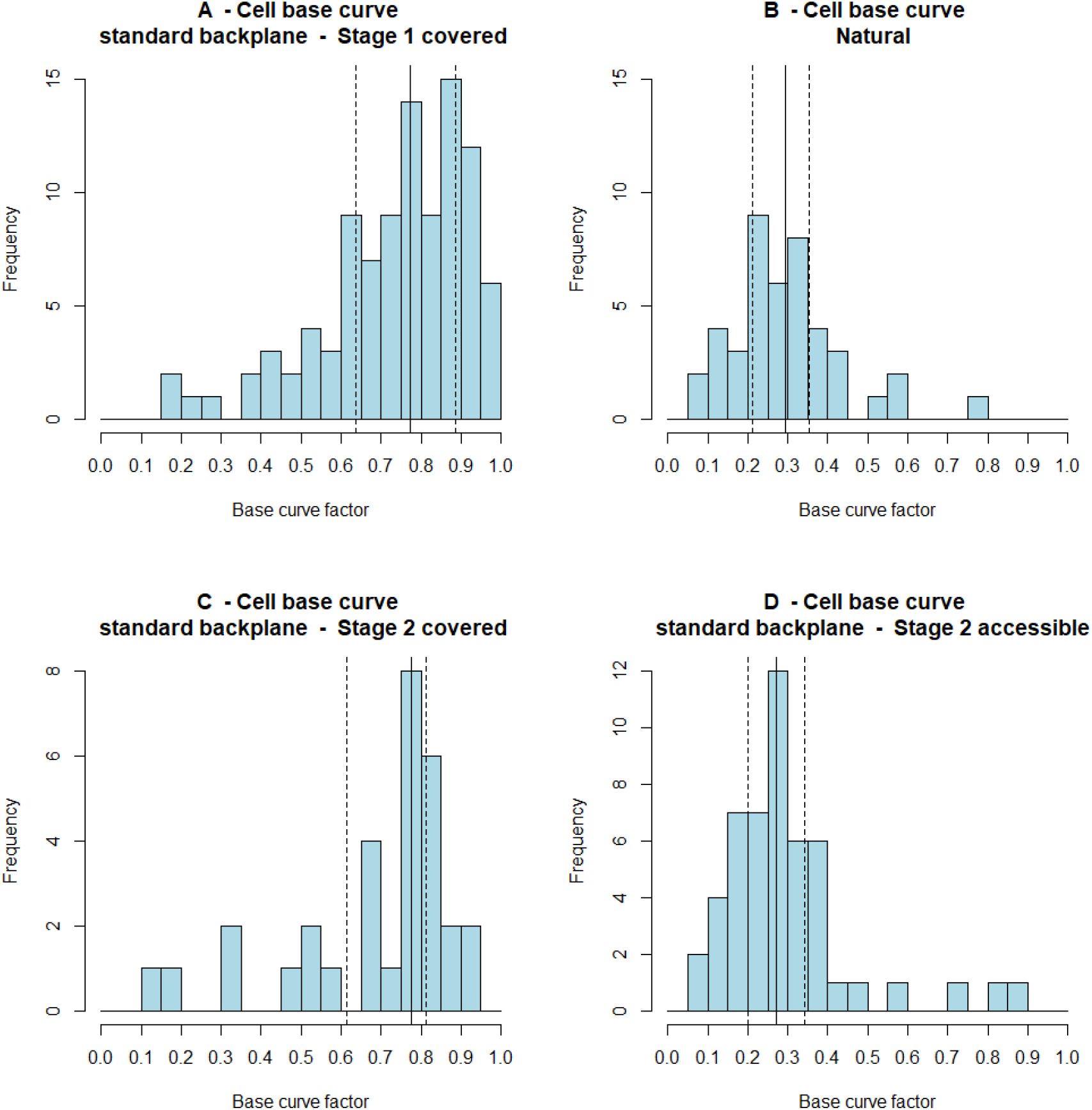
Base curve factor (curved compared with facetted) of cells built on a substrate where the reverse side was shielded. A: the curve factor of bases where access had been restricted. Solid vertical line shows the median, dashed lines the interquartile range. N=99. B: base curve factor of cells sampled from naturally formed comb. N=43. C: base curve factor of cell walls at stage 2 where the access continued to be restricted. N=31. D: base curve factor of cell walls at stage 2 where the access had been allowed. N=50. Bases built where access to both sides is possible are more facetted.

Bases of cells that remained covered had a curve factor of 0.68 ±0.21 across 31 samples. The bases of cells from which the cover had been removed were significantly less curved (0.30 ±0.16, W = 1372, P <0.00001; Figure 0-12 C & D). This demonstrates that cell bases cease to curve and become facetted in the presence of an adjacent, opposing, cell, as predicted (P4.2) - A wall with a cell to both sides will be straight; either formed straight or became straight - derived from the assumed interaction between the bees attempt to construct both adjacent cells with a rounded, convex profiles.

### Experiment iv – curvature of cell base wall built on a thick wax substrate was influenced by isolation or being shared

We measured 86 instances of cell bases accessible from only one side due to initial wax thickness and a further 43 bases found within natural comb were measured as a control sample. Subsequently we measured the bases of 64 cells from the same area after further construction had thinned the substrate.

The curve factor for the isolated bases was 0.81 ±0.14, which was significantly greater than the factor for natural bases (0.30 ±0.14, W = 3643, P <0.00001; Figure 0-13 A & B). This demonstrates that cell base walls curve where there is no constraint imposed by an adjacent cell, as predicted when one assumes that bees attempt to construct cells with a rounded, convex profiles.

**Figure 0-13.**
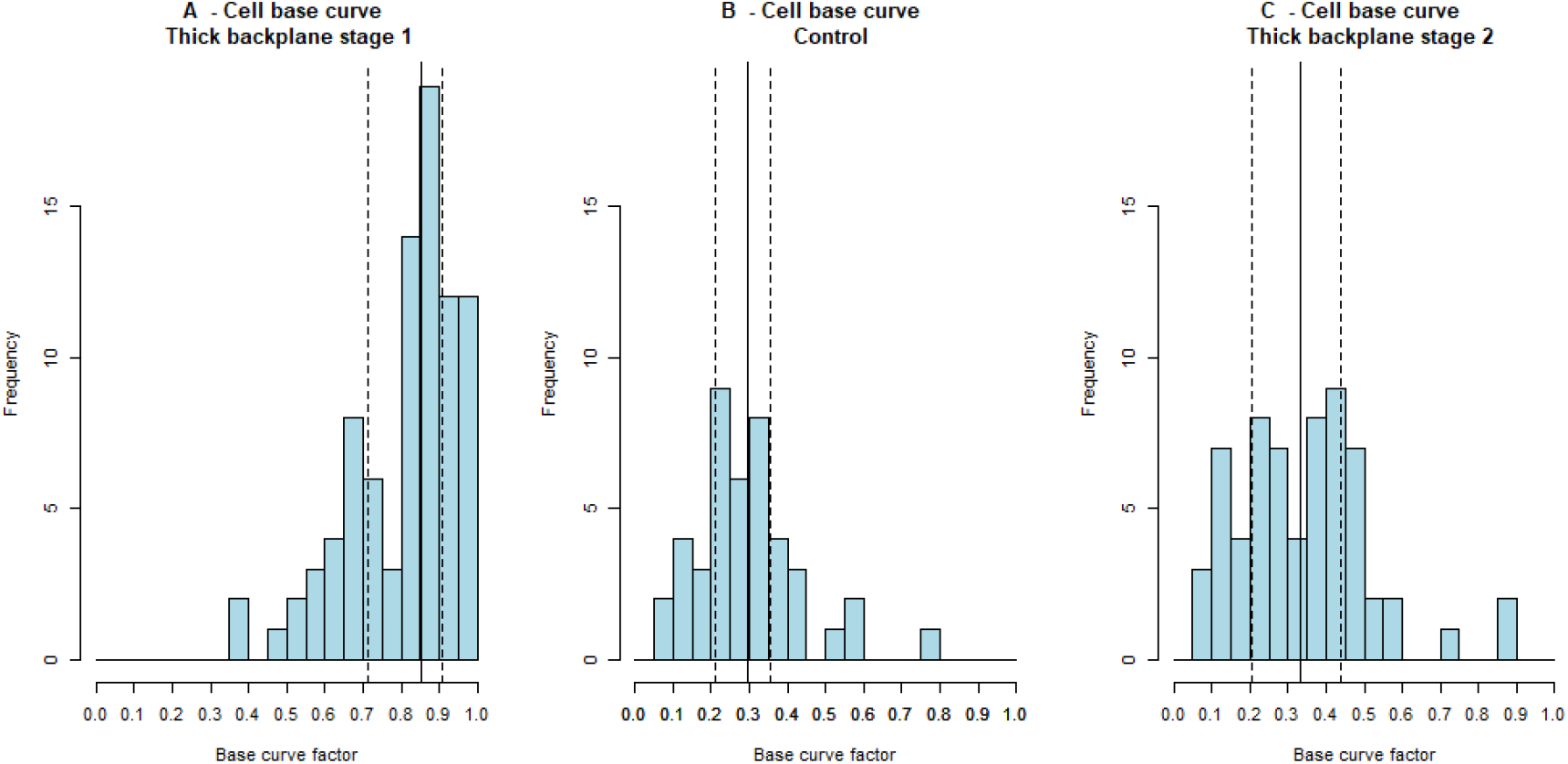
Base curve factor (curved compared with facetted) of cells built on a thick substrate. A: the curve factor of bases to cells at stage 1, depth of 4mm-6mm while base wax was still thick and isolating. Solid vertical line shows the median, dashed lines the interquartile range. N=86. B: base curve factor of cells sampled from naturally formed comb. N=43. C: base curve factor of cell walls at stage 2 where bases had become thin enough for interaction between construction of cells on each side. N=64. Bases built where access to both sides allows interaction are more facetted.

Cell bases at stage 1 had a curve factor of 0.81 ±0.14 across 86 samples. A further 64 measurements taken at stage 2 of experiment showed that the profile of the cell bases had become significantly less curved, more facetted (0.34 ±0.17, W = 5319, P <0.00001; Figure 0-13 A & C). This demonstrates that cell walls are flatter when subjected to influence from both sides, as predicted (P2) - A wall with a cell to both sides will be straight; either formed straight or became straight - derived from the assumed interaction between the bees attempt to construct both adjacent cells with a rounded, convex profiles.

### Base curvature measurement error estimation

A single cell was chosen from the cells formed on the thick backplane samples used in section 0. The chosen cell was measured 24 times with mean = 0.710 Radians, median 0.682 Radians and standard deviation of 0.106 Radians.

The standard deviation, 0.106, was used elsewhere as an estimate of the smallest detectable effect. This value was therefore used as the Smallest Effect Size of Interest (SESOI) when considering equivalence of base curvature between samples.

### Experiment stage 1, isolated walls, and stage 2, shared walls, results compared

Experiments i, ii were used up to stage 1 to test prediction P1 – walls with a cell to just one side would be curved, and stage 2 tested prediction P2 that walls of cojoined cells would be straight. These tests were performed under two conditions, but both considered how the side walls of cells curved when either isolated or shared. The individual results 4i and 4ii have both demonstrated that isolated walls are curved significantly more when isolated and, as can be seen in Figure 0-14 A, the curvature of both forms of isolated walls are similar under both conditions, and both are distinct from the curvature of walls shared by two cells.

**Figure 0-14.**
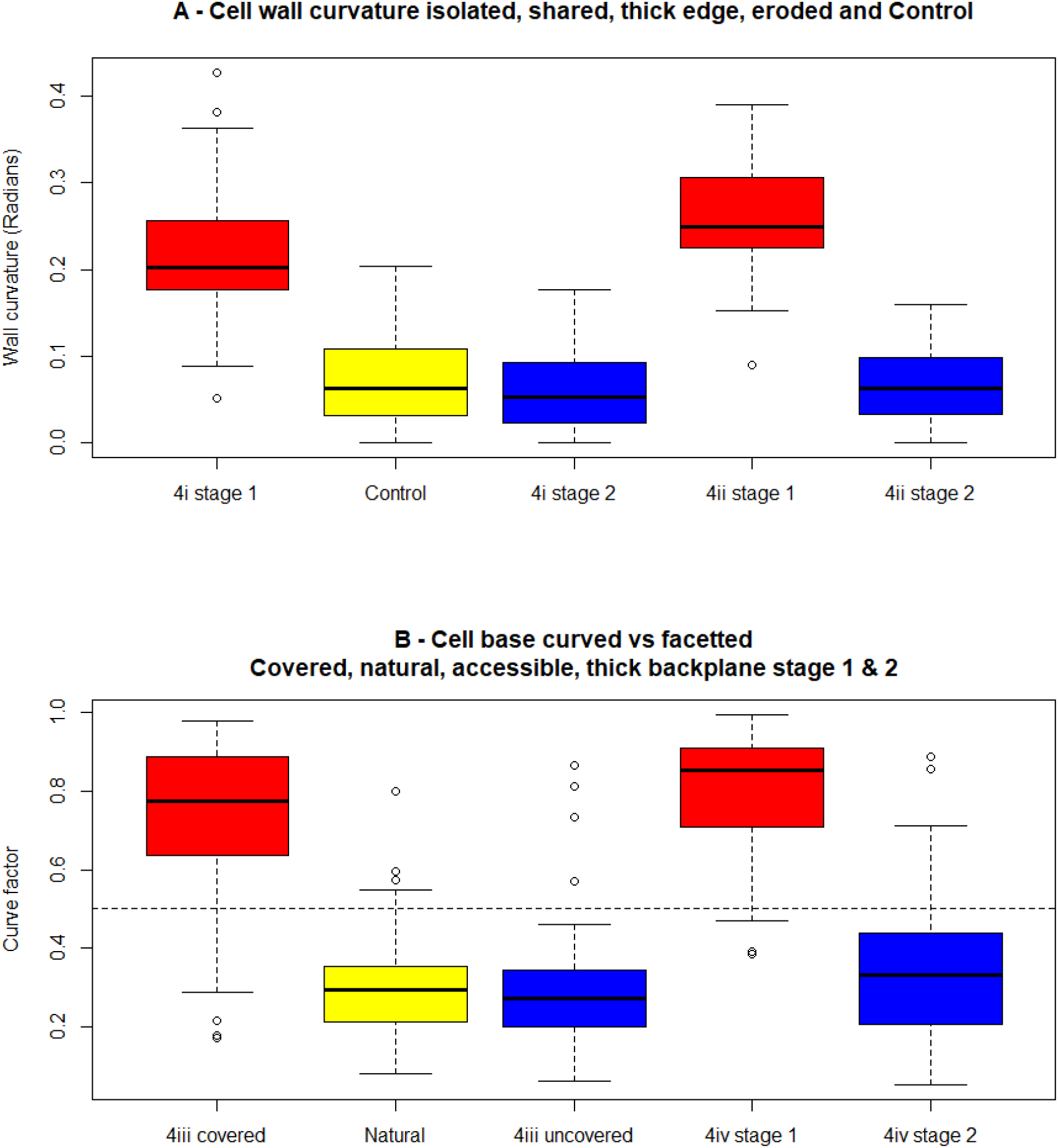
Comparison of wall curvature across all tested conditions. A: curvature of isolated walls, shown in red, isolated by being on the edge of comb (i, N=62) and by being against a thick barrier (ii, N=62), the shared walls used as control are shown in yellow, N= 62. Shared walls, shown in blue, once a second adjacent cell had been built (i stage 2, N= 43) and after the thick barrier had been eroded (ii stage 2, N=43). The curvature of both forms of shared walls and that of natural comb are mutually equivalent. B: curve factor of cell bases. Single-sided access, shown in red, isolated by a barrier (iii, N=99) and by a thick substrate (iv stage 1, N=50): these compared with bases from natural comb shown in yellow, N= 64. Two-sided access, shown in blue, once the barrier had been removed (iii uncovered, N= 31) and after the thick substrate had been eroded (4iv stage 2, N=86). The curvature of both forms of shared bases and that of natural comb are mutually equivalent.

The three categories of shared walls, i stage 2, i control and ii stage 2 were tested for equivalence, each compared with the others hence three pairwise comparisons were performed. Three two-way TOST tests were performed using 0.5 standard deviation as SESOI and in all pairings the walls were equivalently straight (4i stage 2 cf natural P=0.0384 & 0.327, 4ii stage 2 cf. natural P= 0.0112 & 0.876 and 4i stage 2 cf 4ii stage 2 P=0.044 & 0.425).

Experiments iii, iv were used up to stage 1 to test prediction P1, that walls with a cell to just one side would be curved, and stage 2 tested prediction P2 that walls of cojoined cells would be straight. These tests were performed under two conditions, but both considered how the shape of cell bases differed when either isolated or shared. The individual results iii and iv Stage 1 have both demonstrated that isolated bases are curved significantly more when isolated. Also, iii and iv Stage 2 both demonstrated that those bases straightened when the bases became shared. Measurements from all four of those experiments are summarised in Figure 0-14 B.

The shared walls measured in experiment iii stage 2, vi stage 2 and the natural comb were tested for equivalence, each compared with the others hence three pairwise comparisons were performed. Three two-way TOST tests were performed using the measurement error standard deviation as SESOI and in all pairings the walls were equivalently straight (iii stage 2 cf. natural P=0.0009 & 0.941, iv stage 2 cf. natural P= 0.020 & 0.216 and iii stage 2 cf. vi stage 2P=0.024 & 0.189).

Four experiments have been described in this chapter, each of which measured the curvature of a cell wall in a situation where it was subject to builder’s influence from just one side. The four experiments were then allowed to proceed to a second stage where those same cell walls became cojoined to two cells, and therefore subject to influence on both sides. The measurements of all four experiments are summarised in Figure 0-14 sowing the outcome for cell sidewalls in panel A and for cell base walls in panel B. The curvature of walls that were isolated, shown in red, in all four cases differ significantly, and each to a similar extent, from the control cases (shown in yellow). Furthermore, when access to both sides of those walls was possible, the results (shown in blue) were all equivalent to each other and were equivalent to the control cases. Under these experimental conditions, isolated walls were curved, as predicted by my hypothesis III and when circumstances changed the previously curved walls became straight to a degree that the resulting cells were equivalent to naturally formed ones.

## Discussion

Straight walls between each cell are an evident feature of honeycomb, one that has attracted speculation as to how it is built. Stage 1 of these experiments has shown that when building a cell, the bees do not build it as a polygon constructed from flat faces: she builds a convex curved cell. Stage 2 of the experiments studied the effect of additional construction to demonstrate that the cell facets, walls to each side or the base, are flat only because the work undertaken on the other side of that face counteracts that done from within the cell.

Previous attempts to explain the straight faces have been made, proposing thermoplasticity as the mechanism reforming curved walls into flat ones (Pirk et al. 2004; Karihaloo, Zhang, and Wang 2013; Hepburn et al. 2014; Talukdar and Dutta 2019) but having been refuted (Bauer and Bienefeld 2012; Oeder and Schwabe 2017), an alternative explanation is required.

The earlier stage of the experiments has shown that walls with a cell to just one side are significantly more curved than contemporaneous walls with a cell to both sides. This curved nature of isolated walls has been shown for walls at the sides of cells, and for walls on the bases of cells. My experiments at stage 2 went on to show sides and bases of cells are straight when the wall lies between two cells. Experiment 4iii stage 1 caused cell bases to form as domes and after continued construction they remained as domes wherever access was restricted. At the same time, where the barrier had been removed, bases that had been domes were reformed to be facetted. Construction with only single-sided access produces curved cell walls, double-sided access balances the construction effort and so straightens them.

Kepler examined the structure of honeycomb and used a simile of pressing cannonballs together and an example of compaction of pomegranate seeds when discussing how the triple rhombic shape may form at the base of a honeycomb cell (Kepler 1611). Kepler’s description is helpful in forming an image of the interaction between cells and the resulting shape. Darwin also posited that pressing together of deformable cells could lead to the flat sides found on cells in honeycomb (Darwin 1859). Darwin’s conjecture was that precursor species had formed bulbous nest cells and that an intermediate species packed such cells increasingly tightly until the current species of honeybee packed cells closely enough to form hexagons evidence of which Darwin found in earlier literature (Huber 1802).

Cells are not built as discrete items to be pressed together like balloons and at the interface between two cells there is but a single, shared wall. Nonetheless, the descriptions from both Kepler and Darwin illustrate my explanation. The flat nature of the interface between to cells results from a balance of influence exerted on that face by the work undertaken by the builders of each of the cells, one on either side. The first stage of these experiments showed that a cell without external influence will bulge outward leading to a curved cell wall then the results from the later stage of the experiment showed that the presence of a second, balancing, cell will counteract that influence leading to a flat surface. The balance can result from simultaneous construction of both cells in which case the wall will start and remain flat. Alternatively, a wall that is initially curved, having been built in isolation, can be altered once the influence of a second cell is present. Hence, this chapter has shown that the construction of honeycomb is a dynamic system where cell walls are not just fabricated into a fixed, correct, form; rather, the comb, cells and walls are all plastic. During the construction of honeycomb, cell walls are subject to deformation and reformation depending on several interacting influences and it is the eventual balance of these influences that gives rise to straight walls within comb.

## Notes

### Competing Interest Statement

The authors have declared no competing interest.

